# Pupillary dilations of mice performing a vibrotactile discrimination task reflect task engagement and response confidence

**DOI:** 10.1101/444919

**Authors:** DA Ganea, A Bexter, M Guenther, PM Garderes, BM Kampa, F Haiss

## Abstract

Pupillometry, the measure of pupil size and reactivity, has been widely used to assess cognitive processes. As such, changes in pupil size have been shown to correlate with arousal, locomotion, cortical state and decision-making processes. In addition, pupillary responses have been linked to the activity of neuromodulatory systems that modulate attention and perception as the noradrenergic and cholinergic systems. Due to the extent of processes reflected by the pupil, we aimed at resolving pupillary responses in context of behavioral state and task performance while recording pupillary transients of mice performing a vibrotactile two-alternative forced choice task (2-AFC). We show that pre-stimulus pupil size differentiates between states of disengagement from task performance versus active engagement. In addition, when actively engaged, post-stimulus, pupillary dilations for correct responses are larger than for error responses with this difference reflecting response confidence. Importantly, in a delayed 2-AFC task version, we show that even though pupillary transients mainly reflect motor output or reward anticipation following the response of the animal, they also reflect animal decision confidence prior to its response. Finally, in a condition of passive engagement, when stimulus has no task relevance with reward provided automatically, pupillary dilations reflect stimulation and reward being reduced relative to a state of active engagement explained by shifts of attention from task variables. Our results provide further evidence for how pupillary dilations reflect cognitive processes in a task relevant context, showing that the pupil reflects response confidence and baseline pupil size encodes attentiveness rather than general arousal.

**Significance Statement:** For the last 60 years, pupillometry has been used to study various cognitive processes. Among which are mental load, arousal and various decision related components, linking pupil dilations to underlying neuromodulatory systems. Our results provide extensive evidence that in addition to reflecting attentiveness under task performance, pupil dilations also reflect the confidence of the subject in his ensuing response. This confidence coding is overlaid within a more pronounced pupil dilation that reflects motor output or other post-decision components such that are related to the response itself but not to the decision. Our results also provide evidence how different behavioral states, imposed by task demands, modulate what the pupil is reflecting, presumably showing what the underlying cognitive network is coding for.

## Introduction

Pupillometry has been widely used to assess cognitive processes. When observed under constant light conditions, changes in pupil size are reflecting underlying brain activity, presumably mainly as a proxy to Locus Coeruleus (LC) processing (Aston-Jones and Cohen, 2005; Murphy et al., 2014a; Reimer et al., 2016). Though there is also evidence that links pupillary dilations to Colliculi and Cingulate cortex activity, these occur at an increased latency to LC (Joshi et al., 2016). In rodents the cholinergic system has also been shown to correlate with pupillary dilations (Reimer et al., 2016). Such changes in pupil size have been shown to reflect emotional arousal and alertness (Hess and Polt, 1960; Bradley et al., 2008; Vinck et al., 2015), correlate with bouts of locomotion (McGinley et al., 2015; Mineault et al., 2016, Shimaoka et al., 2018), and correlate with synchronized cortical activity (Reimer et al., 2014). Pupil size is also indicative of optimal performance (McGinley et al., 2015; Schriver et al., 2018) since it is taken as a proxy of arousal states that modulate cortical activity and signal processing involved in decision making in rodents (Mineault et al., 2016; McGinley et al., 2015) and humans (Murphy et al., 2014b) exhibiting a U-shaped relationship between baseline pupil size and performance levels. This U-shaped relationship has also been proposed for LC tonic firing levels (Aston-Jones et al., 1999; Usher et al., 1999). This correlates with tonic and phasic LC activity and the LC-NE theory of adaptive gain (Aston-Jones & Cohen, 2005). A tonic dominated LC state would result in overall small or larger pupil size, unresponsive to task events, while phasic LC state would result in lower baseline pupil size and dilations that reflects task relevant events (Aston-Jones et al., 1994; Aston-Jones et al., 1999; Usher et al., 1999; Clayton et al., 2004). Pupil dilations are also a marker of perceptual selection or states of attention switching, indicating as to what underlying cognitive substrate is being perceived (Einhäuser et al., 2008). In addition, when human subjects are actively engaged in a task, such changes in pupil size correlate with an increase in mental effort and cognitive load (Hess and Polt, 1964; Kahneman and Beatty, 1966; Kahneman and Beatty, 1967; Beatty, 1982b) and reflect decision related processes (Preuschoff et al., 2011; Fiedler and Glöckner, 2012; Kloosterman et al., 2015; de Gee et al., 2017) with the decision related component shown to hold information regarding the choice that ends the decision process (Einhäuser et al., 2010) but also decision related information prior to the decision related response (de Gee et al., 2014). Since pupillary responses were shown to occur in response to a variety of behaviors, attention states and overall cognitive function, we aimed at further resolving pupillary dilations during task related behavior in mice. Combining a vibrotactile two-alternative forced choice task (2-AFC) together with pupillometry allowed us to monitor pupillary dilations in context of specific behavioral states as reflected by the degree of task engagement, and levels of task performance as a function of varying difficulty.

Our results show that when subjects are actively engaged in the performance of a task, arousal levels do not influence performance. In addition, pupillary dilations show two distinct epochs. A pre-response phase being a marker of response confidence, continuously reflecting confidence until response time and varying with task difficulty. And a post-response phase, exhibiting a marked dilation relative to the pre-response component that is locked to the response and mainly reflects the motor component of the response or possible reward anticipation.

## Methods

### Animals and Surgery

For all experiments, male C57BL/6J mice were used (Charles River). Experiments were approved by North Rhein-Westphalia State Agency for Nature, Environment and Consumer Protection (Landesamt fur Natur, Umwelt und Verbraucherschutz Nordrhein-Westfalen, LANUV) and conformed to ethical regulations of German Law for Protection of Animal Welfare. For surgery, mice were anesthetized with isoflurane in oxygen (3% induction, 1.5% maintenance; V/V) and body temperature was maintained at 37°C with a feedback-controlled heating pad. Analgesia (Buprenorphine; 0.1mg/Kg) was injected S.C. The fur over the skull was removed and the skin was incised using a scalpel. Several drops of a local analgesia agent (Bupivacaine; 0.25%; Actavis New Jersey, United States) were used for the incision area. Connective tissue was removed and a bonding agent (DE Healthcare Products) was applied over the bone and polymerized with blue light. Next, blue light polymerizing dental cement (DE Healthcare Products) was used to attach a titanium head bar to the skull. Finally, the skin was sutured around the dental cement cap. An antibacterial ointment (Gentamicin) was applied over the surgery area and antibiotics were added to the drinking water of the animals (Baytril; 25mg/ml). Animals were monitored and allowed a week of recovery before training commenced, with food and water *ad libitum*. Mice were housed separately and maintained under an inverted 12 hours light cycle regime.

### Behavior procedure and setup

Mice were trained to perform a vibrotactile 2-AFC task (Mayrhofer et al. 2013). Briefly, upon commencement of training mice were subjected to a water deprivation regime during weekdays, receiving 1 ml of water per training day and water *ad libitum* during weekends. Weight was monitored daily throughout the water deprivation period. If a loss of over 20% body weight was observed compared to the non-deprived weekend days, water was supplemented. Mice were handled and acclimatized to the experimenter for one week. After acclimatization, mice were head fixed for increasing periods of time until accepting 1ml of water while head fixed. Once mice attained this stage, they were placed in the setup on a wheel to monitor locomotion and behavioral training on the detection task began. In general, for the 2-AFC task mice had to detect a target stimulus (90Hz) from two simultaneous bilateral frequencies (for detection distractor was 0Hz, and later for discrimination 10, 20, 40 or 60Hz). Whisker stimuli consisted of one second long repetitive pulses (single-period 120Hz cosine wave) with a maximum deflection amplitude of 400μm. Stimulation frequency was modulated by changing inter-pulse time intervals. Target was randomly delivered to the left or right C1 whisker, stimulated with a piezo bending actuator (Johnson Matthey, Royston, UK) amplified by a piezo controller (MDT693A; Thorlabs, USA) with mice having to report the side the target was presented on by licking on one of two corresponding left or right water spouts placed in front of them (Fig 1A). Lick detection was conducted by capacitive water spouts connected to an Arduino platform (Arduino UNO Rev3; Arduino, Italy). Water delivery was controlled by solenoid valves (Bürkert, Ingelfingen, Germany). Responses to the task were classified under four categories: correct response with mice being rewarded a water drop delivered through the corresponding spout, error response where no water was rewarded, miss when the animal did not respond with a lick within the decision period window, with no water being rewarded or a double-lick when mice licked both spouts within a 60ms period with no water being rewarded. The temporal structure of each trial consisted of a 1 second stimulus presented 1.5 seconds following trials start, with a response window of 2 seconds after stimulus initiation. Inter-trial interval was set as 2 seconds after the response of the animal or end of decision period with a 50% maximal temporal jitter (Fig 1B). Once mice attained a performance of 85% correct responses per session in the detection task, discrimination training commenced. For the *delayed response detection task* water spout movement was controlled by servo electric motors (Savöx, Taiwan), following a determined delayed period after stimulus initiation (1000, 1500, 2000ms). For *passive engagement task*, highly trained mice (>90% correct responses for detection task) were provided with the same target whisker stimulation, as in the detection task, coupled with the automated provision of the reward or in an additional set of experiments provided only with a reward. Control of the behavioral sessions and behavioral data analysis were conducted with custom written LabVIEW (National Instruments, RRID:SCR_014325) and MATLAB software (MathWorks, RRID:SCR_001622).

**Figure 1.**
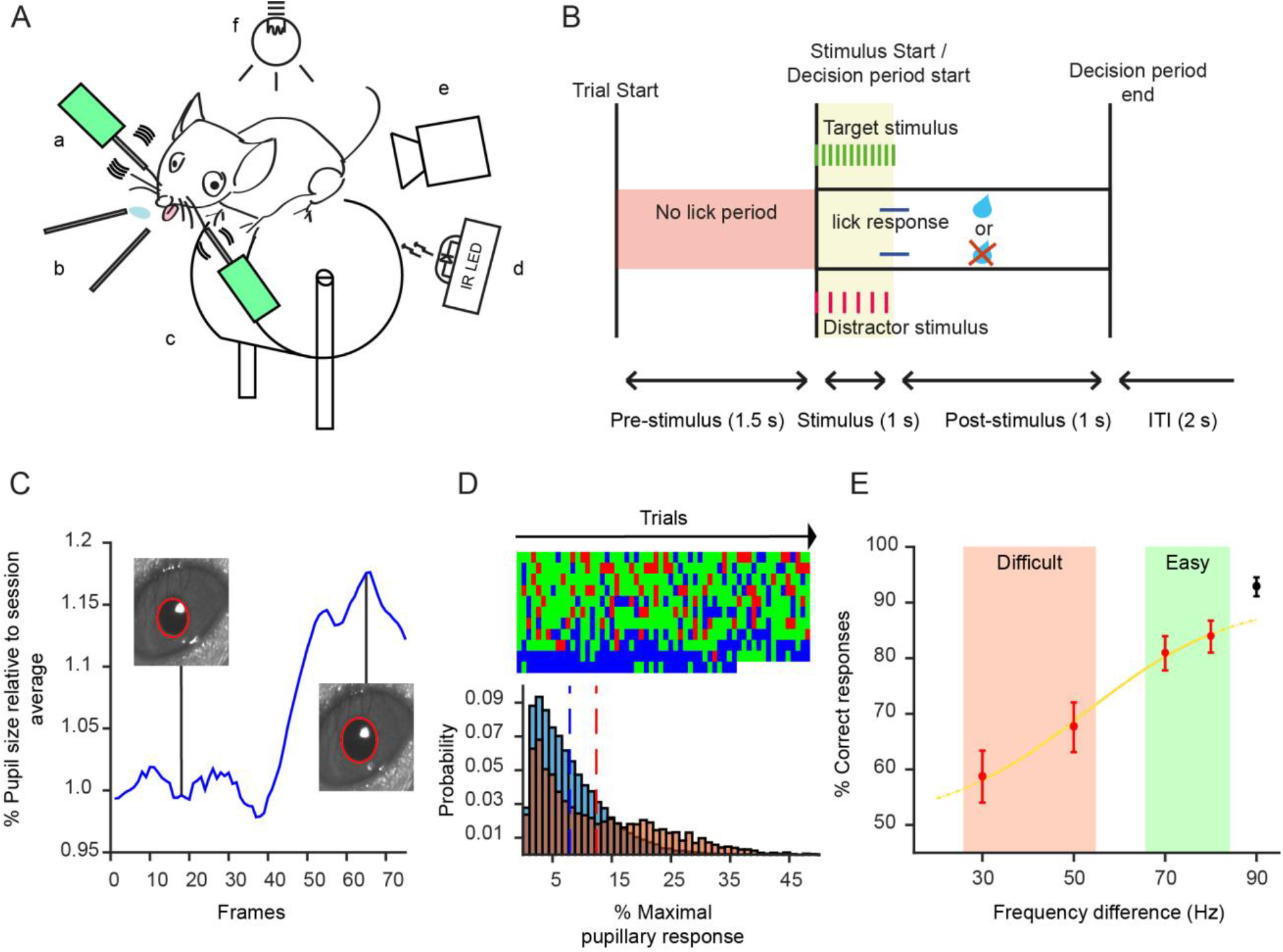
2-AFC task and pupillometry overview. (A) Experimental setup. a - whisker stimulators; b - water spouts; c - wheel; d - IR LED illumination; e - pupil tracking camera; f – ambient illumination. (B) Schematic of a trial sequence for the 2-AFC task used to test animal behavior. (C) Example of a pupillary dilation trace for one behavioral trial. Pupillary trace shown in blue. Insets: examples of pupil detection for two different frames. Recording duration 2.5 seconds. (D) *top* - Example of animal responses (correct - green, error - red, miss - blue) during the performance of the 2-AFC task for a single session. Each row represents 60 trials; *bottom* - Distribution of maximal pupil dilation for non-running (blue) and running trials (red), dashed lines represent distribution mean. (E) Example psychometric response curve for one animal.

### Locomotion

To monitor for locomotion, mice were placed on a Polystyrene (Styrodur®) wheel, 20cm diameter, and movement was tracked using an optical incremental encoder (Optical miniature encoder 2400; Kübler, Germany). Locomotion was determined as movement >5 cm/sec during the duration of the trial.

### Pupil imaging and detection

Images were acquired using a Point Grey Chameleon3 camera (Point Grey Research) at 30 FPS with a 50mm lens and the pupil illuminated by an IR led. Throughout the behavioral session, the setup was maintained under constant white light illumination, with the pupil in a dynamic range. Pupil movies were recorded separately for each trial (15 frames for baseline). For image acquisition, a custom written LabVIEW software (National Instruments, RRID:SCR_014325) was used and pupil detection and fitting was conducted offline with custom written MATLAB software (MathWorks, RRID:SCR_001622). For pupil detection a threshold was determined for each frame and the image converted to a binary image. The pupil was detected using a circle fitting algorithm that detects the mean [x,y] coordinates of the pupil in the binary image. For determining the validity of the detection, 20% random frames in each movie were visually analysed by the experimenter. The validity criterion was set as >98% fit for all non-blinking frames per session. As blinking results in a quick and sudden change in measured pupil size, a threshold for the differential of the pupil transient was used and trials where blinking was detected were removed from all subsequent analysis.

### Data analysis

#### Behavioral data analysis

Psychophysical response curves for each animal were analysed with a MATLAB tool box for psychophysical data analysis (psignifit version 2.5.6; see http://bootstrap-software.org/psignifit, which implements a maximum-likelihood method (Wichmann and Hill, 2001). We used a logistic function

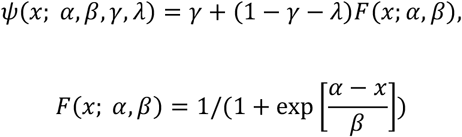

to fit the data points (parameters: *α,β,γ=* 0.5,*λ*[0 0.2]) and obtain the inflection point of the discrimination threshold and slopes. Confidence intervals to the response for each stimulus pair were computed based on a binomial distribution with a confidence level of 95%. Performance in the 2-AFC task was computed as:

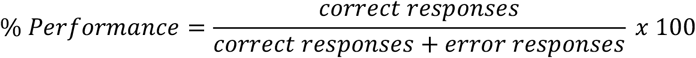

For figure 2B including miss condition performance was computed as:

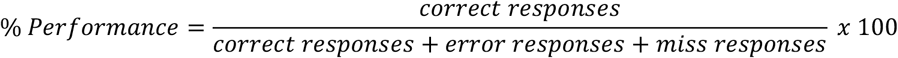

All data used in this study consists of mice having a performance above 85% correct responses per session in the detection task.

#### Pupil data analysis

For comparing pupil dilations between animals and sessions, pupil diameter per data point in each session was divided by the average pupil size of that session, i.e. normalized pupil size. For determining the pupillary dilation transient per trial (change relative to pre-stimulus period), for each trial, the average pre-stimulus, normalized pupil size was calculated and subtracted from each normalized pupil size sample point, with the result divided by the average value of the pre-stimulus normalized pupil size. For quantifying the pupillary response, the maximal pupil dilation per trial was used, resulting in the maximal pupillary response.

#### Experimental design and statistical analysis

The experimental design for baseline period analysis (Fig 2) and post-stimulus analysis (Fig 3) consisted of 8 mice. Delayed response task (Fig 4) consisted of 3 mice and passive engagement task (Fig 5) consisted of 3 mice. For determining significance between the different conditions in the baseline period, pupil size was averaged per trial for the pre-stimulus period, referred to as average pupil size modulation, and a one-way ANOVA used across animals followed by a Tukey post hoc (multiple comparison) test. For determining significance between conditions of the post-stimulus pupillary response, the maximal pupil size following stimulus onset was used as a test variable, and this maximal pupillary response analysed using a one-way ANOVA across animals followed by a Tukey post hoc (multiple comparison) test. For correlations we used a one-sided Kendall rank coefficient for test statistic and a linear fit of the data was applied. Unless stated otherwise shaded error bars for the pupillary dilation transient represent 95% Confidence Interval (95CI) and error bars represent Standard Error of the Mean (SEM). Data in text is presented as mean±SEM. Statistical analysis was conducted using MATLAB software (MathWorks, RRID:SCR_001622).

**Figure 2.**
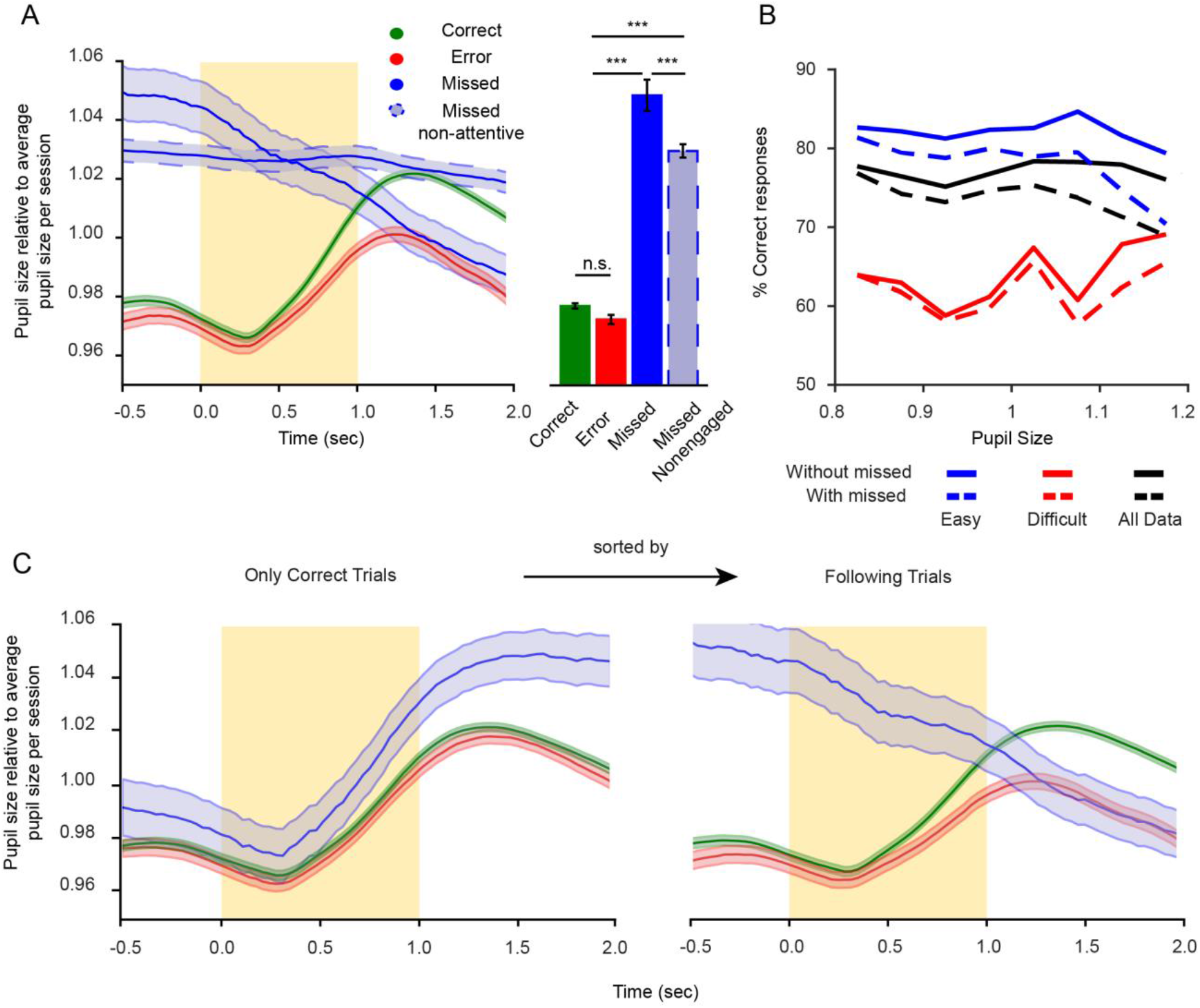
Pre-stimulus pupillary size of mice performing a 2-AFC task reflects task engagement state and holds no information regarding subsequent performance. (A) *left*: Average pupil size for all trials separated into correct trials (green), error trials (red), miss trials (solid-blue) and miss trials during the non-attentive period at the end of the session (dashed-blue). *right*: Average pupil size for pre-stimulus period. (B) Performance for all animals (n=8) in dependence of baseline pupil size. Performance shown for three categories: all trials (black), difficult trials (distractor > 30Hz; red), easy trials (distractor ≤ 30Hz; blue) and under two conditions: miss trials included (dotted lines), miss trials excluded (solid lines). (C) Dilation transients showing the history dependency of the pupil size using an example of rewarded trials and trials following rewarded trials only. *left*: Only rewarded (correct) trials are shown. The trials are separated by the outcome of the following trial. *right*: Trials following rewarded (correct) trials. Yellow rectangle represents stimulus.

**Figure 3.**
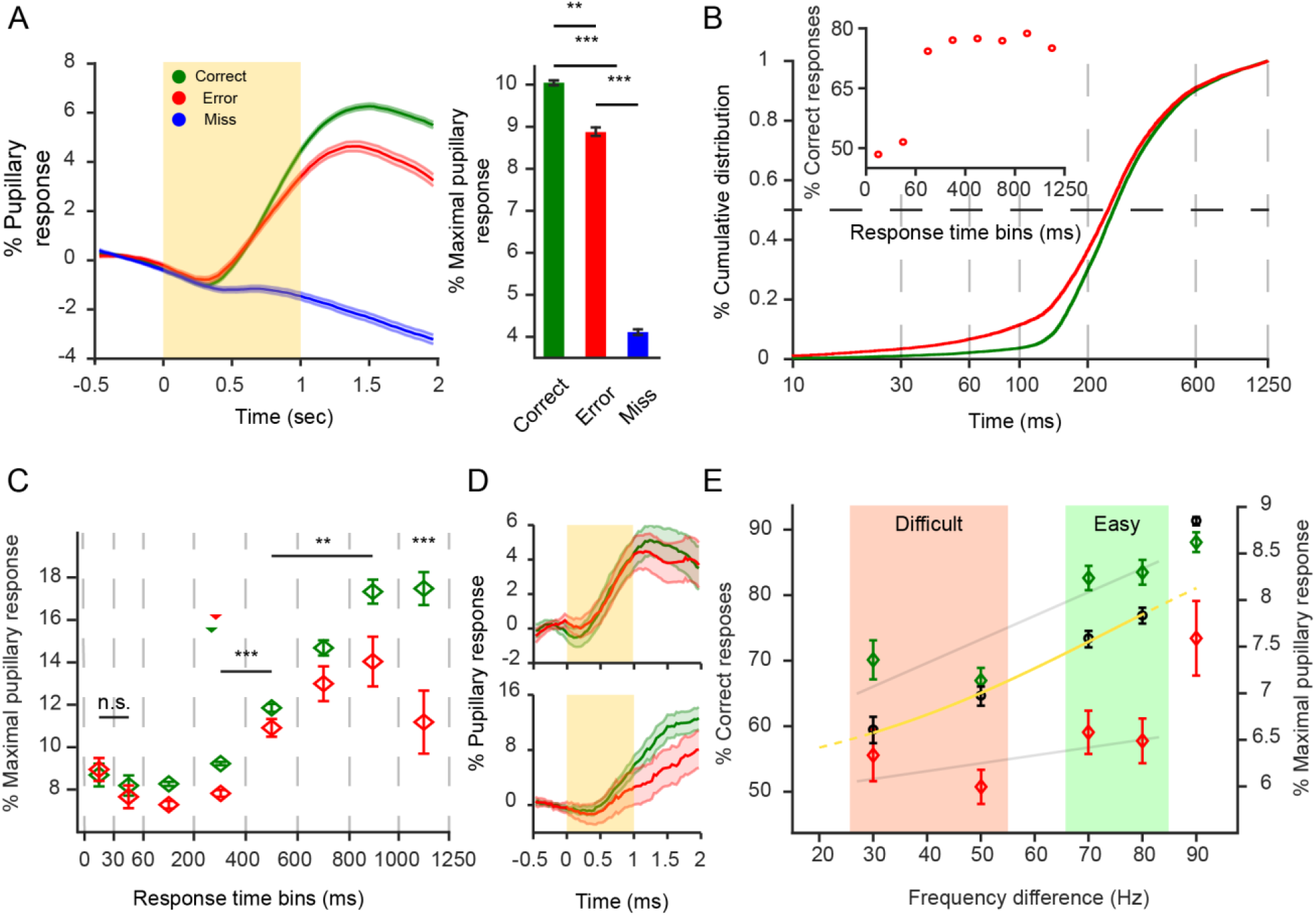
Pupillary dilation transients of mice performing a 2-AFC task differ depending on animal response and reflect internal decision components. (A) *left* - Pupil dilation transients for mice performing a 2-AFC task (n=8 mice; N=92 sessions) normalized relative to baseline period for correct (green), error (red) and miss (blue) responses. *right* - Pupil response magnitude following stimulus onset for the different response types. (B) Cumulative distribution function (CDF) plot comparing response time distributions for correct (green) and error (red) responses. inset - %correct responses as a function of RT bin. (C) Pupil response magnitude for different RT time bins for correct (green) and error (red) responses. Dashed vertical lines represent time bins used for averaging pupil response magnitude. Average response time represented by triangles for correct (green - 264ms) and error (red - 260ms) responses. (D) Example average pupillary dilation traces for correct (green) and error (red) responses in two response time bins. *top* - 0-30ms RT bin. *bottom* - 1000-1250ms RT bin. (E) Decrease in pupil response magnitude correlates with decrease in animal performance as difficulty increases for performance in a 2-AFC discrimination task for correct (green) responses but not for error (red) responses. Black circles are average performance across mice with logistic fit (yellow). Grey lines represent linear fit for discrimination task. Yellow rectangle represents stimulus.

**Figure 4.**
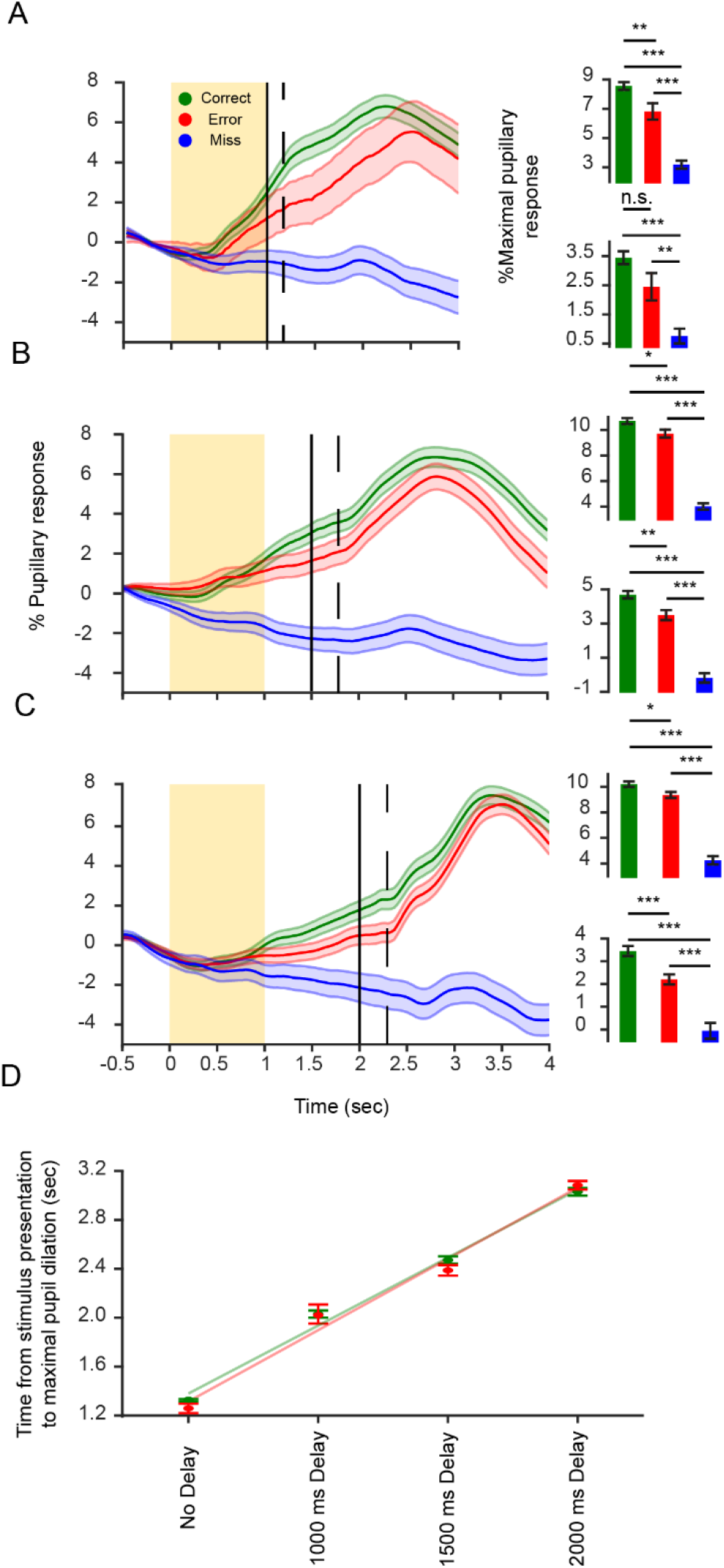
Pupillary transients in a delayed response 2-AFC detection task reflect decision variables but mainly encode for post decision components. (A-C) *left in each panel* - Average pupillary dilation transients for correct (green) error (red) and miss (blue) responses in a delayed response detection task (n=3 mice). Spout delay from stimulus onset 1000ms (7 sessions); 1500ms (8 sessions); 2000ms (9 sessions); respectively. Yellow rectangle represents stimulus; Black vertical line represents water spouts presentation; Dashed vertical line represents mean RT. *bottom right in each panel* - Average percent maximal pupillary response in a period of 500ms before water spout presentation. *top right in each panel* - Average percent maximal pupillary response in the period following water spout presentation. (D) Time difference between stimulus onset and maximal pupillary response increases as a correlation of the delay period for correct (green) and error (red) responses. Green and red lines represent linear fit for correct and error responses respectively.

## Results

### Pre-stimulus pupillary size of mice performing a 2-AFC task reflects task engagement state and holds no information regarding subsequent performance

The pupil of mice was tracked while performing a 2-AFC task (Fig 1A and B). Pupil size was observed to fluctuate throughout the session and through each individual trial (Fig 1C). It is known that locomotion correlates with an increase in pupil size (McGinley et al. 2015; Vinck et al. 2015; Mineault et al. 2016). Our data also shows this phenomenon, with pupillary responses being dominated by larger dilations when the animal is locomotive (Fig 1D). As we wanted to examine pupillary dilations in respect to task performance, all locomotion corresponding trials were removed from analysis to avoid contamination of the pupillary response to the task by locomotive and hyper arousal (McGinley et al. 2015; Neske et al., 2018) states. When performing a 2-AFC task there are three different possible behavioral response types: correct, error and miss (no response) with correct and error categorizing behavior into a task engaged state and miss trials indicating a task disengaged state. Mice showed a high degree of task engagement, represented by their consistent response to the presented stimuli throughout each session (84.1±4.9% of all trials), receiving a reward for correct responses or no reward for error responses (Fig 1D). However, the miss condition could be separated into two different types of behaviors, sparse miss responses during extensive periods of engagement (attentive period) or as a batch at the end of the session (non-attentive period). It is possible for these separate miss responses to have a different pupillary phenotype. As such a cutoff criterion was used when 50% of the trials within a 10 trial window consisted as miss. Miss trials before this cutoff were categorized as being during an attentive period and miss trials following this were categorized as being during a non-attentive period. In addition, the 2-AFC task enabled us to separate performance for different stimuli in easy and difficult task categories while tracking pupil dilations for these categories (Fig 1E). In the period prior to stimulus onset (baseline period) pupillary dilation traces revealed a significant difference between the four response conditions for the average baseline pupil dilation trace (Fig 2A; F(3,20060)=321.74, p<0.001), with mice that are in a state of disengagement from task performance having a baseline pupil size larger than when in the engaged states with no difference in baseline pupillary size for correct and error responses (M_(correct)_=0.977±0.001; M_(error)_=0.972±0.002) but a significant difference for the attentive and non-attentive miss conditions (M_(attentive miss)_=1.048±0.001; M_(non-attentive miss)_=1.029±0.002). The observed difference in both baseline size and pupillary transient between the attentive and non-attentive miss responses might arise due to a history dependence of the dilations. Indeed, in the non-attentive period the pupil is constantly enlarged during repeated miss trials (Fig. 1D and 2B). However, in the attentive period, miss trials occur rarely (Fig 1D). To analyze the history of increased pupil size during miss trials we looked into the trials preceding a miss trial (Fig 2C). To reduce the effect of different trial conditions on pupil size, we restricted our analysis to correct trials preceding a miss trial.

However, similar results were observed when restricting to miss after error trials (data not shown). We found, in the attentive period, that miss trials occur with fast switches in pupil size starting at the end of the previous trial and reaching average baseline levels again already after a single miss trial (Fig. 2C). In order to test whether the baseline period holds information regarding task performance, animal performance was analyzed in respect to baseline pupil size (Fig 2B) restricted to the attentive period. When baseline pupil size is observed as a function of performance over all trials including miss response trials, performance is constant for small and medium baseline pupil size but drops for above average pupil size, resulting in a significant negative correlation (Fig 2B, black dashed line; r_τ_=-0.57, p=0.03). However, this effect might be mediated solely by miss trials which have a larger baseline pupil size overall. Indeed, when performance is observed solely for the engaged state excluding miss trials, it does not drop as a function of baseline pupil size (Fig 2B, black solid line; r_τ_=0.07, p=0.64). As such, baseline pupillary size seems to hold no perceptual information regarding task performance (engaged state) or optimal task performance. In addition, we observed no effect of task difficulty over this phenotype when tested for a negative correlation for easy tasks or positive correlation for hard tasks (Fig 2B, blue and red; r_τ easy all data_=- 0.57, p=0.03; r_τ easy without miss_ =0.21, p=0.27; r_τ hard all data_= 0.07, p=0.27; r_τ hard without miss_=0.36, p=0.14), with performance levels dropping overall per difficulty level but remaining constant as a function of baseline pupil size.

### Pupillary dilation transients of mice performing a 2-AFC task differ depending on animal response and reflect internal decision components

Thus, to observe perceptually related task responses as reflected by the pupil, the pupillary dilation transient was baselined relative to the pre-stimulus period, resulting in a pupillary dilation transient that reflects the perceptual content of the information withheld by the pupil. We observed a significant difference between pupillary dilation transients for the three different response types (Fig 3A; F(2,19396)=1272.84, p<0.001). Pupil dilation transients for the disengaged state remained principally unchanged following whisker stimulation and revealed only a late (~700ms) and barely noticeable pupillary response (M_(miss)=_4.06±0.08). Contrary to this, pupillary dilation transients for the engaged state, showed a faster (~330ms) and increased response following stimulation onset, for both correct (M_(correct)_=9.96±0.05) and error (M_(error)_=8.79±0.09) with correct responses showing a larger pupillary response magnitude than error responses (Fig 3A). This observed difference between the pupillary dilation transients for correct and error responses might simply arise due to coding of externally induced signals such as reward attainment for the correct condition or possibly from internal decision components such as confidence coding as for a subject highly trained in a task, correct responses would be supported by more confidence. As response time (RT) is indicative of the decision we analyzed the underlying dependency of the RT distribution, because this might reflect varying confidence in the response or decisions which might differ between very early responses to late responses. RT distribution analysis for correct and error responses (Fig 3B) shows that the two distributions diverge from one another for early responses (<120ms), with relative error density being more prominent in this interval and behavior being at chance levels. In addition, response accuracy in the early bins (<60ms) indicates that early RTs are responses not guided by stimulus information (Fig 3B). Hence, we wanted to observe whether the pupillary response diverges between correct and error responses as a function of when the RT is provided, by observing the pupillary response magnitude in different RT bins. Analysis revealed a significant difference between pupillary response magnitude for correct and error responses per different RT bins (Fig 3C; F(15,18068)=112.43, p<0.001). However, there is no difference between the pupillary response magnitude of correct and error responses for very early responses (see also figure 3D top), when provided in the first 60 milliseconds (M_(correct,0-30ms)_=8.68±0.53; M_(error,0-30ms)_=8.95±0.55; M_(correct,30-60ms)_=8.19±0.52; M_(error,30-60ms)_=7.66±0.57). Importantly, this shows that the observed difference for the pupillary dilation transient between correct and error responses is not due purely to the difference in reward attainment, motor output or reward anticipation, hence rather reflecting internal decision variables. For RT bins where the choice is guided by the stimulus (i.e performance above chance levels) (60-1250ms) the pupillary response magnitude increases as a function of the increase in RT with pupil response magnitude being larger for correct responses than for error throughout all bins (M_(correct,60-200ms)_=8.27±0.10; M_(error,60-200ms)_=7.29±0.17; M_(correct,200- 400ms)_=9.22±0.08; M_(error,200-400ms)_=7.82±0.16; M_(correct,400-600ms)_=11.85±0.17; M_(error,400-600ms)_=10.91±0.37; M_(correct,600- 800ms)_=14.68±0.29; M_(error,600-800ms)_=12.99±0.65). For late RT bins (>800ms) this phenotype is altered, with the pupillary response magnitude for correct responses continuing to increase while the response magnitude for error trials not increasing further (see also figure 3D bottom) (M_(correct,800-1000ms)_=17.33±0.41; M_(error,800-1000ms)_=14.04±0.84; M_(correct,1000- 1250ms)_=17.48±0.46; M_(error,1000-1250ms)_=11.18±0.89). In addition to any RT underlying influences on the pupillary transients, we hypothesized that there might also be a task difficulty effect. As the difference between target stimulus and distractor stimulus decreases, it becomes more difficult to discriminate between the two simultaneously presented stimuli and solve the task. Indeed, this effect is seen in the average psychometric response curve for all mice (Fig 3E). Performance was highest for the detection task (M=91.3±0.6% correct responses) and dropped with increasing the frequency of the distractor, reaching near chance levels (M=59.4±2.0% correct responses). This enabled us to observe pupillary response magnitude relative to task difficulty as experienced by the animals. This decrease in performance correlates with a decrease in the pupillary response magnitude for correct responses but not with error responses which show no significant decrease as a function of difficulty level for the discrimination task (Fig 3E; r_τ correct_=0.388, p<0.001; r_τ error_=0.003, p=0.402). Hence, as performance drops to chance levels the pupillary response magnitude for correct and error trials tends to converge, indicating that when choice is random pupillary dilation becomes similar. Taken together, this further suggests that pupil dilation codes for the response confidence of the animal.

### Pupillary transients in a delayed response 2-AFC detection task reflect decision variables but mainly encode for post decision components

Due to the slow kinetics of the pupillary response, it is possible that any perceptual response reflected by the pupil in the period between presentation of the stimulus and RT would not be observed due to task design (temporal separation between stimulus and RT). In order to determine if such pupillary perceptual representations of the task do occur during this period, we tracked the pupil of mice performing a delayed response detection task with the water spouts being presented after a delay period following stimulus onset. Pupillary dilation traces for both correct and error responses began increasing following stimulation (Fig 4 A-C), showing a significant difference between the pupillary dilation trace for correct, error and miss responses already before the animal provided its response to the task or receives any response feedback (F(2,968)_1000ms_=25.06, p<0.001; F(2,2247)_1500ms_=76.66, p<0.001; F(2,2279)_2000ms_=32.08, p<0.001). This increased pupillary dilation trace for correct and error responses was maintained throughout the stimulus – RT interval (1000ms: M_correct_=3.45±0.22; M_error_=2.45±0.47; M_miss_=0.76±0.25; 1500ms: M_correct_=4.69±0.22; M_error_=3.49±0.29; M_miss_=-0.18±0.27; 2000ms: M_correct_=3.45±0.22; M_error_=2.20±0.22; M_miss_=-0.06±0.35). Following the RT of the animal, pupillary responses exhibited a second and more pronounced, significant increase in pupillary dilation (F(2,968)_1000ms_=70.73, p<0.001; F(2,2247)_1500ms_=139.52, p<0.001; F(2,2279)_2000ms_=91.68, p<0.001) with correct trials still showing the largest pupil dilation and miss trials hardly any difference (1000ms: M_correct_=8.56±0.27; M_error_=6.82±0.56; M_miss_=3.18±0.27; 1500ms: M_correct_=10.69±0.23; M_error_=9.71±0.31; M_miss_=4.02±0.25; 2000ms: M_correct_=10.20±0.22; M_error_=9.35±0.23; M_miss_=4.25±0.32). Average pupillary dilation trace for the disengaged state remained overall unchanged as in the un-delayed task, indicating the unresponsiveness of the pupil when mice are disengaged. For the engaged state, across all delay periods, the increase in delay and the pupillary response magnitude exhibited a positive correlation for both correct and error responses (Fig 4D; r_τ correct_=0.541, p<0.001; r_τ error_=0.436, p<0.001), indicating that the maximal pupillary _İ_ dilation follows the RT and not the stimulation. For all delay periods, baseline pupillary size was larger for the disengaged state (miss condition) versus the active engagement state as in the un-delayed task, with baseline for correct and error not being significantly different (data not shown). The difference in pupillary dilation transients observed for correct and error responses in the stimulus-RT interval (Fig 4A-C) reflects a stimulus-based decision prior to RT with a second component due to motor output or possible reward anticipation.

### Pupillary transients of mice in a state of passive engagement hold perceptual information for both stimulus and reward and reflect a state of quasi-engagement

To better determine how and if reward and stimulus presentation contribute to pupillary responses we conducted two additional sets of experiments. In the first type of experiment, mice already trained in the 2-AFC task, were provided automatically with a water reward upon whisker stimulation (90 vs. 0Hz) in all trials for several sessions. In the second type of experiment, the same mice were now only provided with a water reward, without whisker stimulation, for several sessions, in order to observe whether there is a pupillary representation of the stimulation in addition to the reward. For both experiments the temporal sequence of the task was the same as in figure 1B. Hence, in both cases mice were passively engaged in the task, meaning they were responding to the presented reward without the requirement to solve a task to obtain it so that the whisker stimulation loses its task relevant meaning. When comparing the passive engagement states with correct responses provided in an active engagement state (as all three conditions contain the attainment of reward) there is a significant difference in pupillary response magnitudes between both passive engagement states and active engagement (Fig 5A; F(2,4788)=177.58,p<0.001). For passive engagement, the presentation of the *stimulus + reward* versus *reward only* elicited a significantly higher pupillary response magnitude (M_(reward)_=4.35±0.16; M_(reward + stimulus)_=5.03±0.16). This indicates that both stimulus and reward *per se* are encoded by the pupillary response. However, the pupillary response magnitude is higher when mice are actively engaged as in the *stimulus + reward* condition (M_(active engagement)_=7.94±0.13). Within passive engagement states there was a significant difference between engaged states, when mice responded to the reward and disengaged states, when mice did not respond to the reward (miss condition) (Fig 5B; F(3,2704)=20.29, p<0.001) with baseline pupillary size increased when mice were in a state of disengagement versus engagement*, stimulus + reward* condition (M_engaged stimulus + reward_=0.985±0.003; M_disengaged stimulus + reward_= 1.06±0.02) and *reward only* condition (M_engaged reward_=0.991±0.003; M_disengaged reward_=1.032±0.007). Interestingly, when analyzing the pupillary response magnitude in different RT bins, there is a significant difference between the groups (Fig 5C; F(5,2034)=49.41, p<0.001). For early RTs (<60ms) the pupillary response magnitude for both passive engagement conditions is not significantly different (M_reward_=5.41±0.44; M_reward + stimulus_=5.90±0.49) but for responses provided around the median RT (*reward + stimulus*,117ms; *reward only*,159ms) for both conditions, the pupillary response magnitude is higher for *reward + stimulus* (M_reward_=4.63±0.24; M_reward + stimulus_=5.73±0.25), indicating the stimulus being perceived in the response decision period. For late RTs (200 – 400ms) the pupillary response magnitude is dropping sharply for both conditions and there is again no significant difference between the conditions (M_reward_=1.47±0.27; M_reward + stimulus_=1.65±0.25). This unresponsiveness of the pupil to task occurrences can be explained by observing baseline pupil size (Fig 5E) which is significantly different across time bins for both the *reward only* condition (F(2,1028)=15.84, p<0.001) and *reward + stimulus* condition (F(2,1006)=18.5, p<0.001). Baseline pupil size remains small and is not significantly different for early and median RTs (M_reward 0-60ms_=0.974±0.008; M_reward 60-200ms_=0.976±0.004; M_reward + stimulus 0-60ms_=0.960±0.008; M_reward + stimulus 60-200ms_=0.972±0.004), indicating a state of engagement, but increases sharply for late RTs (M_reward 200-400ms_= 1.011±0.005; M_reward + stimulus 200-400ms_=1.003±0.004), explaining the observed drop in the pupillary response magnitude, with the pupil becoming unresponsive to task occurrences as baseline pupil size increases due to a state of disengagement. Hence, the same task variables are differentially reflected by the pupil following varying behavioral requirements imposed on the animal.

**Figure 5.**
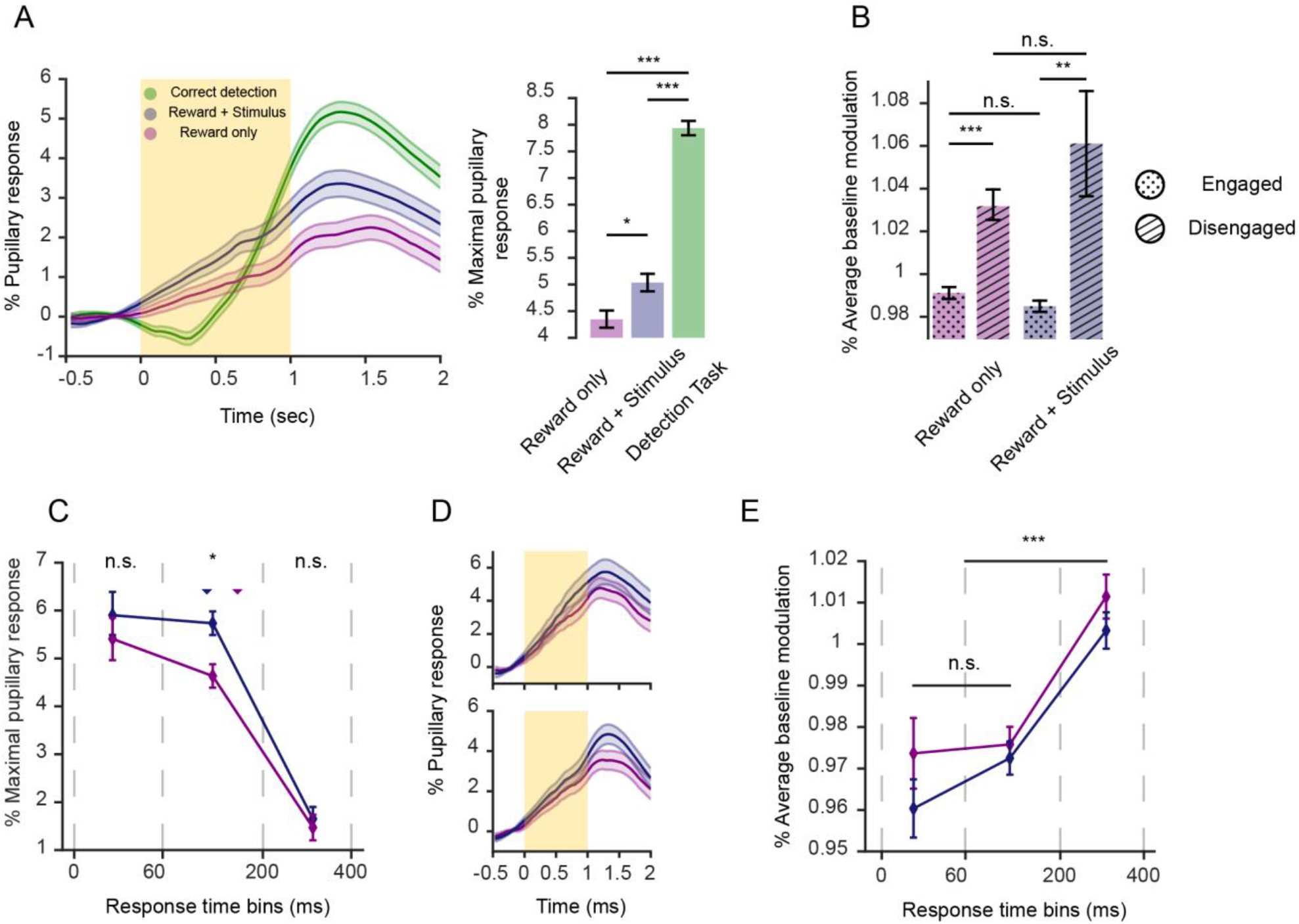
Pupillary transients for a state of passive performance hold perceptual information for both stimulus and reward reflecting a state of quasi-engagement. (A) Pupil dilation transients differ for states of active performance versus passive performance. *left* - Averaged transient of percent change in pupil size relative to baseline period, before stimulus onset/reward delivery, (n=3 mice) for reward only (purple; 1311 trials), reward + stimulus (blue; 1123 trials) and correct responses in a detection task (green; 2419 trials). *right* - Average percent maximal pupil response following stimulus onset/reward delivery for the different behavioral situations. (B) Average pupil size during baseline period before stimulus onset relative to average pupil size per session for the passive performance states and their corresponding disengagement periods. (C) Pupil response magnitude for different RT time bins for reward only (purple) and reward + stimulus (blue). Triangles represent average RT for reward + stimulus (blue - 117ms) and reward only (purple - 159ms). (D)*top* - Pupillary dilation transients for RTs in the 0-60ms bin. *bottom* - pupillary dilation transients for RTs in the 60-200ms bin. (E) Maximal pupillary response as percent change from baseline for different RT bins shown for the passive performance states reward only (purple) and reward + stimulus (blue). Vertical dashed lines show behaviorally relevant RT bins. Yellow rectangle represents stimulus.

## Discussion

Under conditions of task engagement, pupil dilations in humans were shown to represent various aspects of underlying cognitive activity, among which prediction error (Preuschoff et al., 2011; Braem et al., 2015; Urai et al., 2017), reward anticipation (Chiew and Braver, 2013) and response confidence (Lempert et al., 2015). Studies using mice might provide an advantage in linking the underlying mechanisms of pupil dilations with neuronal activity, however these are still emerging, and it remains unclear to what extent pupil dilations in mice represent the same complex aspects of cognitive function as in humans. In the current study we show that when in a state of engagement, defined as task responsiveness, large dilations are observed following stimulation, with larger dilations for correct than error responses (Fig 3A). However, this difference might be reward driven (Lee and Margolis, 2016). Relating pupil dilation to RT revealed for impulsive, non-stimulus induced responses, (Carpenter and Williams, 1995; Mayrhofer et al., 2012), no significant difference between correct and error dilations and a chance level performance in these trials (Fig 3B and C). Thus, the correct-error dilation difference does not originate from reward related mechanisms *per-se* or from motor responses linked to licking behavior. A reward driven representation would imply a difference that is not RT dependent. As all other external task variables are maintained constant per trial, this implies that the difference would rather reflect a stimulus dependent decision component. Though most pupil studies in humans indicate prediction error representation, our results are not in line with this idea. This would imply increased dilations for errors and a correct-error difference also for early RTs, since non-evidence based prediction should be the same, while reward outcome differs. In addition, any coding for a decision component should be represented in the post-stimulus and pre-RT period. To test for this, we conducted a delayed-response task that temporally separates decision components from motor responses. Two distinctly observed dilation periods were observed (Fig 4A-C). One, a slow dilation following stimulation, and a second, more pronounced dilation after the response. These findings further support decision related effects, as transients already differed in the first, pre-feedback period exhibiting the correct-error difference. The second dilation, locked to RT, relates to motor response or reward anticipation as post decisional variables. Where does this correct-error difference arise from? It is possible to explain it in terms of a representation of response confidence, a notion which our findings support. First, confidence coding is supported by the early RT results, as response confidence would be equal or irrelevant when responses are random and not evidence based. Second, the correct-error difference was observed only when responses were stimulus based, with dilation transients continuing to increase with increased RT (Fig 3C). Indeed, for trained animals, responses leading to a correct outcome should be accompanied by higher choice confidence. The correct-error difference in dilation increases as a function of RT and indicates that response confidence is maintained by the mechanisms underlying pupillary dilations until RT, awaiting feedback. Third, response confidence related pupillary dilations were also reflected by differences in dilations in respect to task difficulty. Higher response confidence being exhibited as larger dilation for easier tasks but dropping with increased difficulty (Fig 3E) and as difficulty increases, leading to performance dropping to chance levels, and the correct-error difference converging. When guessing, response confidence would be equal. Fourth, in support of confidence coding, a recent study (Lak et al., 2017) linked response confidence with the dopaminergic system. Indeed, LC modulates dopaminergic activity in both Ventral Tegmental Area (VTA) and Substantia Nigra (Grenhoff et al., 1993; Zhu, 2018) and VTA afferents innervate LC (Ornstein et al., 1987). It is conceivable that the dopaminergic system reflects confidence through dilations either driven by LC activation of dopaminergic loci or prefrontal cortex feedback arising from these interconnected systems (Arnsten and Goldman-Rakic, 1984; Sara and Hervé, 1995; Jodo et al., 1998).

Pupil studies in mice, mainly studied pupil size in correlation with arousal levels (Reimer et al., 2014; Murphy et al., 2015a) indicated through locomotion (Mineault et al., 2016; Shimaoka et al., 2018) or surprise (Vinck et al., 2015). Further, arousal levels influence performance (McGinley et al., 2015; Schriver et al., 2018) manifested as a U-shaped relationship between the two (Murphy et al., 2011). Though, see Kahneman and Beatty, 1967; Beatty, 1982a; Karatekin et al., 2007; Neske and McCormick, 2018. We show that under task performance, baseline pupil size is not directly an arousal marker but rather indicates engagement. Pre-stimulus pupil size is smaller when engaged versus disengaged from task performance, as defined by a lack of task responsiveness and manifested by a lack of pupil reactivity to stimulation, contrary to dilations under task engagement (Fig 2A). The switch between engagement and disengagement can occur quickly from trial to trial (Fig 2C). Overall, when disengaged, the pupil is not overtly coding task relevant occurrences. In the present task, we show that performance was higher for small and mid-range pupil sizes but dropped for larger sizes that tend to relate with disengagement. Indeed, when excluding miss trials performance remained steady (Fig 2B). Indicating the performance drop for larger pupil sizes results from miss responses not varying arousal. The discrepancy between previous results (McGinley et al., 2015) and ours, might be due to task modality, different cognitive requirements imposed by the 2-AFC task and Go/noGo task response categorization, where perceptual failure or lack of motivation are less distinguishable. Under low light conditions used in previous studies, pupil size might be mainly influenced by sympathetic input. However, parasympathetic input would dominate in the ambient light condition used in the present study (Steinhauer et al., 2004). Hence, we conclude that baseline pupil size, holds no information for perceptually relevant task processing in the presented behavioral scenario. Pre-stimulus pupil size rather reflects engagement or disengagement states, not a general state of arousal.

The behavioral state or task demands, based on what various underlying neuronal mechanisms are directed towards, may well influence pupil diameter. As such, stimulus-response association was altered by decoupling the stimulus detection requirement from reward attainment. A condition we refer to as passive performance. Under these conditions, dilations were smaller compared to correct responses under active performance, even though stimulus and reward presentation were experienced the same (Fig 5A). In the passive state, when both reward and stimulation were presented, dilation was increased compared to when only reward was presented, indicating that under passive performance both reward and stimulus are reflected by the pupil. It has been previously reported that LC neurons exhibit a small early component which is stimulus related (Rajkowski et al., 2004). Thus, when actively performing the task, pupil dilations are dominated by internal decision variables related to response confidence, while under passive performance external occurrences dominate the dilation. Further, contrary to active performance, where pupil size increases together with RT (Fig 5C), an opposite relationship is observed between pupil size and RT under passive performance. This is explained by engagement state, which has an opposite trend to the pupillary dilation (Fig 5E). Increased baseline pupil size and no pupil reactivity to task occurrences for late RTs, even though mice still responded to reward, show within trial disengagement that may be related to fluctuations of attention. The possibility of baseline pupil size to reflect task disengagement or within trial attention switches is also manifested by the observed history dependence of pupil dilations for miss condition during attentive periods (Fig 2C). Hence, under the passive performance state an underlying state of attention switching is reflected. This may relate to the LC adaptive gain theory and the exploration-exploitation modes (Aston-Jones and Cohen, 2005; Aston-Jones et al., 1994; Aston-Jones et al., 1999; Usher et al., 1999; Clayton et al., 2004). Hence, when the stimulus has no task function, attention quickly fluctuates from an exploitative mode, reflected by low baseline and pupillary reactivity to relevant occurrences, to an explorative mode where task occurrences are not coded, exhibited by increased baseline and low pupillary reactivity to the occurrence. This indicates that while mice are still responsive, they already transition into a state of quasi-disengagement with baseline pupil size reflecting task attentiveness rather than mere arousal. Thus, the behavioral state and requirements posed by the environment are determining what the pupil reflects.

Taken together, our results provide further evidence for the complexity of what pupillary dilations reflect and support findings related to the LC-NE adaptive gain theory. When actively performing the task, these dilations reflect task relevant decision representations of response confidence. Also, baseline pupil size reflects states of task engagement or attentiveness rather than general arousal. Finally, in a state of passive performance, pupillary dilations reflect external occurrences with animal state fluctuating between engaged and disengaged states, relating to fast fluctuation in attentiveness. The presented paradigm combined with pupillometry promises to provide a valuable framework to relate behavioral states with large scale neuronal network dynamics recoded using multielectrode (Jun et al., 2017) or two-photon imaging (Margolis et al. 2014; Stirman et al., 2016) techniques during perceptual decision making.

